# Solagigasbacteria: Lone genomic giants among the uncultured bacterial phyla

**DOI:** 10.1101/176263

**Authors:** Eric D. Becraft, Tanja Woyke, Jessica Jarett, Natalia Ivanova, Filipa Godoy Vitorino, Nicole Poulton, Julia M. Brown, Joseph Brown, C.Y.M. Lau, Tullis Onstott, Jonathan A. Eisen, Duane Moser, Ramunas Stepanauskas

## Abstract

Recent advances in single-cell genomic and metagenomic techniques have facilitated the discovery of numerous previously unknown, deep branches of the tree of life that lack cultured representatives. Many of these candidate phyla are composed of microorganisms with minimalistic, streamlined genomes lacking some core metabolic pathways, which may contribute to their resistance to growth in pure culture. Here we analyzed single-cell genomes and metagenome bins to show that the “Candidate phylum SPAM” represents an interesting exception, by having large genomes (6-8 Mbps), high GC content (66%-71%), and the potential for a versatile, mixotrophic metabolism. We also observed an unusually high genomic heterogeneity among individual SPAM cells in the studied samples. These features may have contributed to the limited recovery of sequences of this candidate phylum in prior metagenomic studies. Based on these observations, we propose renaming SPAM to “Candidate phylum Solagigasbacteria”. Current evidence suggests that Solagigasbacteria are distributed globally in diverse terrestrial ecosystems, including soils, the rhizosphere, volcanic mud, oil wells, aquifers and the deep subsurface, with no reports from marine environments to date.

## INTRODUCTION

Technological innovations in single-cell genomics and metagenomics have led to a rapid improvement in our understanding of the genomic features, evolutionary histories and metabolic capabilities of tens of phylum-level branches of Archaea, Bacteria and Eukarya that lack cultured representatives (Yoon et al., 2011; Rinke et al., 2013; Becraft et al., 2015; Brown et al., 2015; Castelle et al., 2015). In these efforts, the subsurface has emerged as a bountiful reservoir of undiscovered, deeply branching microbial lineages that may hold clues to the emergence and evolution of life on our planet (Kallmeyer et al., 2012; Colwell and D’Hondt, 2013). Many of the recently discovered candidate phyla are composed of microorganisms with streamlined genomes lacking some core metabolic pathways, which may be a factor contributing to the inability to obtain pure cultures of these organisms (Rinke et al., 2013; Becraft et al., 2015; Brown et al., 2015; Castelle et al., 2015). This apparent genomic reduction has given rise to hypotheses of genome-streamlining, parasitism, symbiotic lifestyles, and large-scale community metabolic inter-dependence (Giovannoni et al., 2014; Castelle et al., 2015; Anantharaman et al., 2016).

Our preliminary findings from several subsurface environments indicated that the ‘Candidate phylum Spring Alpine Meadow’ (SPAM) constitute an intriguing exception to genome streamlining in oligotrophic environments. The existence of this lineage was first suggested by several 16S rRNA gene sequences obtained in 2004 from an alpine soil from the Colorado Rocky mountains (Lipson, 2004). Subsequently, related 16S rRNA gene sequences were identified on all continents except for Antarctica in environments such as crop soils (Hansel et al., 2008; Chen et al., 2012; Figuerola et al., 2015), copper mine soil (Rodrigues et al., 2014), the subsurface oxic sediments of Hanford Formation at Pacific Northwest National Laboratory (PNNL) (Lin et al., 2012), subsurface groundwater from the Rifle site (Anantharaman et al., 2016), as well as volcanic mud and oil wells (unpublished) (Figure 1).

**Figure 1.**
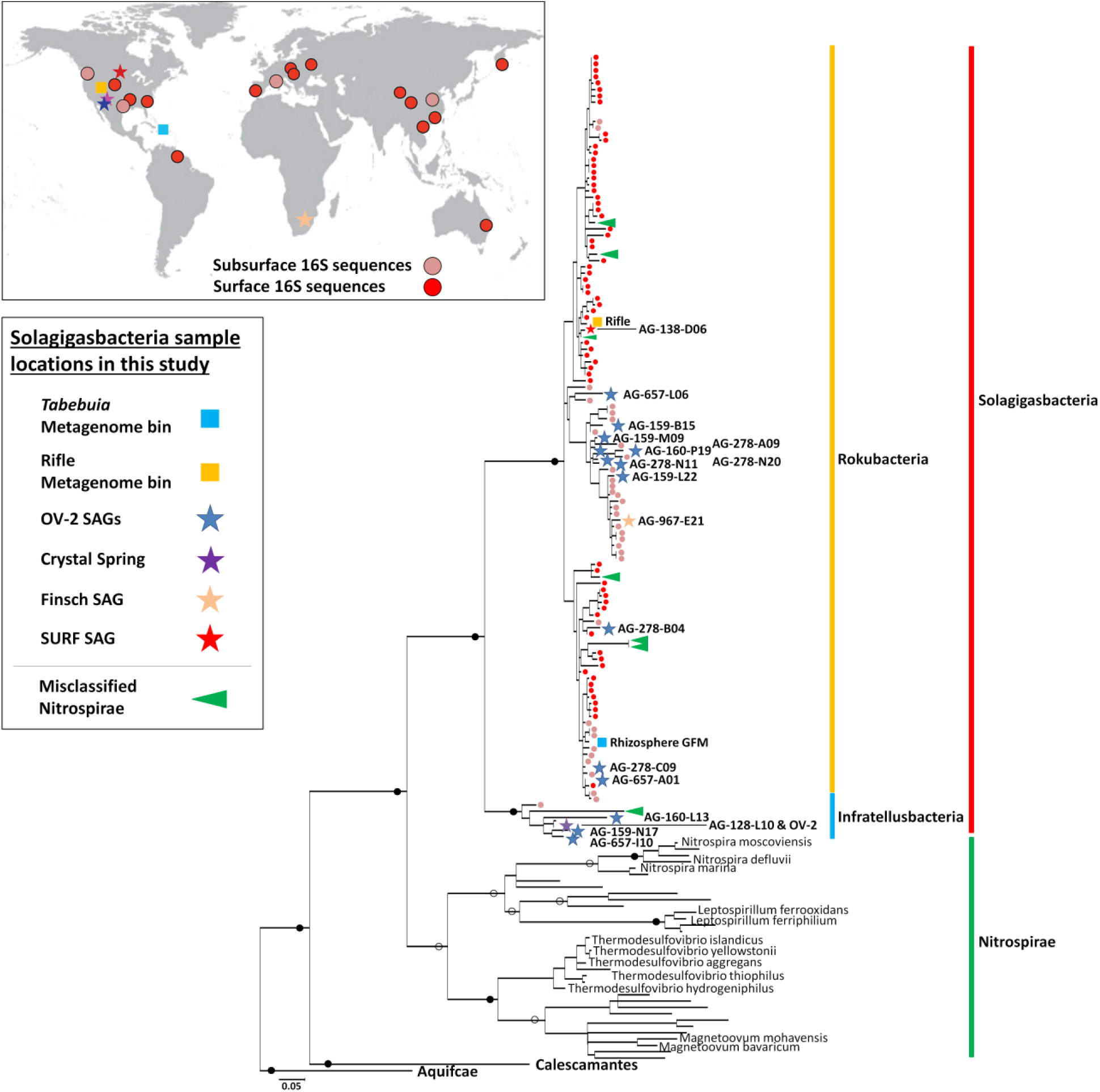
Maximum-likelihood phylogeny of the Solagigasbacteria (red line), based on partial 16S rRNA gene sequences (∼600 bps in length). Included are NCBI sequences with ≥ 85 % nucleotide identity to Solagigasbacteria SAGs. The Rokubacteria are demarcated by a dark orange line, and the Infratellusbacteria are demarcated by a light blue line. Nitrospirae genome 16S rRNA gene sequences representing classified genera and sequences misclassified as Nitrospirae are demarcated by a green vertical bar and green arrows, respectively. 16S rRNA gene sequence from previous surface (red circles), and subsurface (tan circles) studies are also indicated at the terminal branch of each sequence. Map insert (upper left) shows geographic distribution of reported Solagigasbacteria 16S rRNA gene sequences from past surface (red circles), and subsurface (tan circles) studies. Sequence identifiers are reported in Supplemental Figure 3. Previously sampled subsurface sites from Puerto Rico and Nevada where Solagigasbacteria 16S rRNA genes sequences were identified are not shown in Figure 1 insert due to space constants. Stars indicate Solagigasbacteria SAGs (blue = OV-2, purple = Crystal Spring; tan = Finsch mine; and red = SURF) and squares indicate metagenome bins (light blue = *Tabebuia*; orange = Rifle site), and color corresponds to site SAGs were isolated from (left; also see Supplemental Figure 2). All SAGs and metagenome bins that contained a 16S rRNA gene are included in the phylogeny. Full circles indicate bootstrap values >90 %; open circles indicate boot-strap values >70 %. Scale bar represents 0.05 nucleotide substitutions per site.

Here we present genomic sequences from 19 individual SPAM cells from aquifers of different depths in Nevada, South Dakota and South Africa. We compare genomic data from these individual cells to SPAM metagenome bins from Nevada groundwater, a *Tabebuia* rhizosphere in Puerto Rico, and a prior study of the Rifle DOE Scientific Focus Area (SFA) in Colorado (Anantharaman et al., 2016), where the first metagenome bins of these organisms were obtained. This first global 16S rRNA gene survey of SPAM suggests that they comprise a monophyletic, phylum-level lineage that is most closely related to Nitrospirae. Different from the Nitrospirae, SPAM genomes are consistently large, with high %GC and the potential for a mixotrophic metabolism, all packaged within small cells. SPAM cells are also characterized by an unusually high genomic heterogeneity among individuals, with no environments identified to date with near-clonal populations. The unique combination of large genomes encoding the ability for a generalist metabolic strategy in oligotrophic environments, and contained within small cells, is a rare observation among the recent explosion of candidate phyla characterization (Castelle et al., 2015; Anantharaman et al., 2016; Hug et al., 2016b). High level of genetic heterogeneity among SPAM individuals in studied samples is another intriguing feature that may present certain challenges to their investigations.

Based on these observations, we propose renaming SPAM to the candidate phylum “Candidatus Solagigasbacteria” (hereby referred to as Solagigasbacteria), with reference to “sola” and “gigas” (Latin for “lone” and “giant”), encompassing the previously identified Rokubacteria and the newly identified class in this study, the “Candidatus Infratellusbacteria”, with reference to “infra” and “tellus” (Latin for “below” and “Earth”).

## MATERIALS AND METHODS

### Field sample collection

Shallow aquifer water samples were collected from a groundwater evaluation well in Nye CO, Nevada, USA, named “Oasis Valley 2”, hereafter referred to as “OV-2”, on 14 December, 2014 (36.96° N, −116.72° W). OV-2 is a PVC-cased hole, drilled in alluvial sand and gravel derived from tertiary volcanics to a depth of 36.5 m in 2011. The well is screened (i.e. perforations were cut into the casing through which water can enter, but sand and other aquifer materials do not) over the interval from 9.1 – 27.4 m. Samples OV-2 P1, P2 and P3 were collected after removal of one, three and ten well volumes at a pumping rate of ∼10,600 L/min. Microbial biomass was collected on 0.2 μm polyethersulfone membrane filters (Millipore, Sterivex) from one, three, and five liters of samples at time points OV-2 P1, OV-2 P2, OV-2 P3, respectively.

Discharge water samples were collected from the Crystal Spring, which is located adjacent to Death Valley, CA, USA on 13 December, 2014 (36.42° N, −116.72° W). Crystal Spring is the largest spring of the largest Oasis of the Mojave Desert; Ash Meadows, Nye CO, NV, USA. It is located within the discharge zone for a regional aquifer hosted within the highly fractured Paleozoic carbonates in the Death Valley Regional Flow System (DVRFS) (Belcher et al., 2009).

Subsurface water samples were collected from water at the Sanford Underground Research Facility (SURF) at 91.4 meters below land surface (mbls) in Lead, South Dakota on 12 December, 2014 (44.35° N, −103.75 W). SURF samples were collected from perennial wall seeps associated with century-old horizontal legacy drifts in metamorphic rock. Subsurface water samples were also collected from a borehole at Finsch diamond mine at a depth of 857 mbls in South Africa on 11 November, 2012 (-28.38° S, 23.45° E).

All aquatic samples were collected aseptically from flowing pumped lines (OV-2 and Crystal Spring) or directly from the source (SURF and Finsch). For single cell genomics, one-milliliter aliquots were amended with 5% glycerol and 1x TE buffer (all final concentrations), frozen on dry ice in the field and stored at −80°C until further processing. For metagenomics, the DNA from OV-2 samples were extracted from microbial biomass collected on 0.2 μm polyethersulfone membrane filters (Millipore, Sterivex) using the MO BIO PowerSoil DNA Isolation Kit (MO BIO Laboratories Inc., Carlsbad, CA) according to manufacturer’s protocol. An additional freeze/thaw cycle was included after the addition of solution C1 and immediately prior to the ten-minute vortex step (30 min at −80°C followed by 10 min at 65°C). Additionally, a sample for metagenome sequencing was collected from a *Tabebuia* (*T. heterophylla)* rhizosphere in the serpentine area of Cabo Rojo Puerto Rico on 12 March, 2013. Three secondary roots from one tree, about 15-20 cm in length were collected, cut, stored in a 50 mL polyethylene centrifuge tube and transported on ice to the laboratory. The rhizosphere samples were obtained by washing the roots with 25 mL 1X PBS/Tween20 and shaken at 240 rpm horizontally for 1 hour, and frozen at −80 °C. The PBS/Tween20 solution with the rhizosphere was centrifuged at 9,000 g for 20 min at 4 °C. Genomic DNA was extracted from the resulting pellet using the MoBio PowerSoil DNA Isolation Kit with bead tubes (Carlsbad, CA) following Earth Microbiome Project standard protocols (http://www.earthmicrobiome.org/protocols-and-standards/). Site images and the physicochemical characteristics of these field samples are reported in Supplemental Figure 2 and Supplemental Table 1.

**Figure 2.**
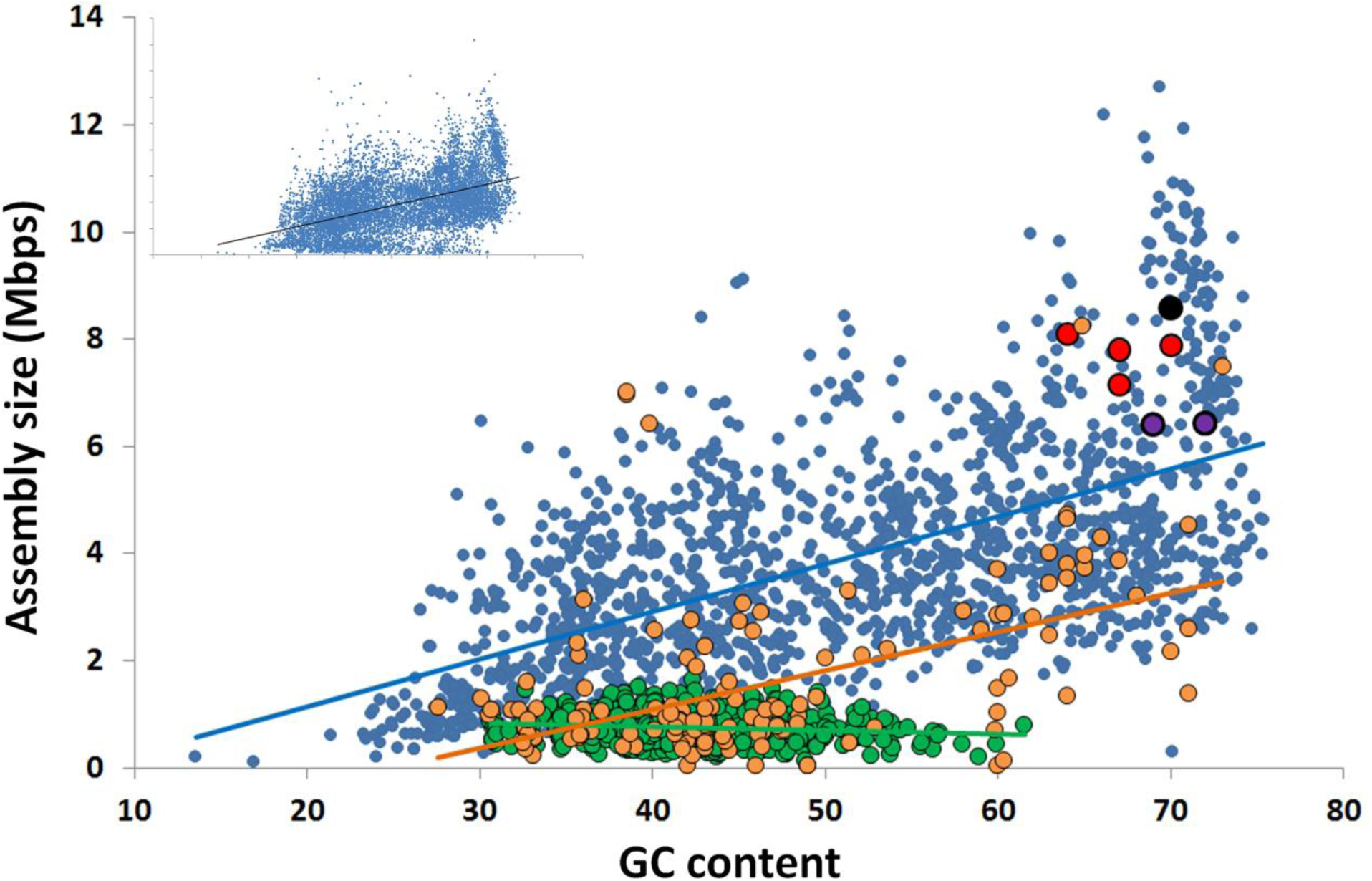
Correlation between the G+C content and genome size among all finished bacterial genomes in IMG (blue; R^2^=0.35), The 4 most complete Solagigasbacteria SAGs from OV-2 (red), metagenome bins from OV-2 (bin 8) and Puerto Rican *Tabebuia* rhizosphere that contain a 16S rRNA gene (purple), and the Rifle metagenome bin that contains a 16S rRNA gene (black) (also see Supplemental Table 3). Also displayed are the estimated genome sizes for Candidate Phyla Radiation (CPR) genomes (green; R^2^=0.02) and candidate phyla genomes not a part of the CPR lineage (orange; R^2^=0.25). The insert contains all genomes in IMG, including all partial SAG and metagenome bins from all bacterial phyla (R^2^=0.35).

### Single-cell genomics

The generation, identification, sequencing and *de novo* assembly of single amplified genomes (SAGs) was performed at the Bigelow Laboratory Single Cell Genomics Center (scgc.bigelow.org). The cryopreserved samples were thawed, pre-screened through a 40 μm mesh size cell strainer (Becton Dickinson) and incubated with the SYTO-9 DNA stain (Thermo Fisher Scientific) for 10-60 min. Fluorescence-activated cell sorting (FACS) was performed using a BD InFlux Mariner flow cytometer equipped with a 488 nm laser and a 70 μm nozzle orifice (Becton Dickinson, formerly Cytopeia). The cytometer was triggered on side scatter, and the “single-1 drop” mode was used for maximal sort purity. The sort gate was defined based on particle green fluorescence (proxy to nucleic acid content), light side scatter (proxy to size), and the ratio of green versus red fluorescence (for improved discrimination of cells from detrital particles). Individual cells were deposited into 384-well plates containing 600 nL per well of 1x TE buffer and stored at −80 °C until further processing. Of the 384 wells, 317 wells were dedicated for single particles, 64 wells were used as negative controls (no droplet deposition), and 3 wells received 10 particles each to serve as positive controls. Index sort data was collected using the BD FACS Sortware software. The DNA for each cell was amplified using WGA-X, as previously described in Stepanauskas et al. (Stepanauskas et al., 2017). Cell diameters were determined using the FACS light forward scatter signal, which was calibrated against cells of microscopy-characterized laboratory cultures (Stepanauskas et al., 2017).

Illumina libraries were created, sequenced and de novo assembled as previously described (Stepanauskas et al., 2017) This workflow was evaluated for assembly errors using three bacterial benchmark cultures with diverse genome complexity and %GC, indicating 60% average genome recovery, no non-target and undefined bases and the following, average frequencies of misassemblies, indels and mismatches per 100 kbp: 1.5, 3.0 and 5.0 (Stepanauskas et al., 2017). checkM v1.0.6 (Parks et al., 2015) was used to calculate estimated the completeness of assemblies of environmental SAGs. We did not co-assemble SAGs due to the high genomic heterogeneity among individual cells. All SAGs were deposited in the Integrated Microbial Genomes database at the Joint Genome Institute (accession numbers pending).

The 16S rRNA gene sequences were aligned using SINA alignment software (Pruesse et al., 2012). Phylogenetic trees were inferred by MEGA 6.0 (Tamura et al., 2013) using the General TimeReversible (GTR) Model, with Gamma distribution with invariable sites (G+I), and 95% partial deletion for 1,000 replicate bootstraps. SAG assemblies were analyzed for protein encoding regions using RAST (http://rast.nmpdr.org/) (Aziz et al., 2008), and genes (protein families) were annotated with Koala (KEGG) (http://www.kegg.jp/ghostkoala/) (Kanehisa et al., 2016) and InterProScan v5 (Jones et al., 2014). Average nucleotide identity (ANI) and average amino acid identity (AAI) of reciprocal hits were calculated using the online tools at the Kostas Lab website Environmental Microbial Genomics Laboratory (http://enveomics.ce.gatech.edu/aai/) (Goris et al., 2007; Rodriguez R and Konstantinidis, 2014). Synteny plots were produced using the Joint Genome Institute Integrated Microbial Genomes (IMG) system (https://img.jgi.doe.gov/) (Markowitz et al., 2014). Phage genes and transposases were identified as in Labonte et al. (Labonte et al., 2015b).

### Metagenomic sequencing and analysis

For OV-2 samples, 1 ng of DNA was fragmented and adapter ligated using the Nextera XT kit (Illumina). The ligated DNA fragments were enriched with 12 cycles of PCR and purified using SPRI beads (Beckman Coulter). For the *Tabebuia* rhizosphere sample, 100 ng of DNA was sheared to 300 bp using the Covaris LE220 and size selected using SPRI beads (Beckman Coulter). The fragments were treated with end-repair, A-tailing, and ligation of Illumina compatible adapters (IDT, Inc) using the KAPA-Illumina library creation kit (KAPA biosystems). For both OV-2 and *Tabebuia* rhizosphere metagenomes, qPCR was used to determine the concentration of the libraries, and libraries were sequenced on an Illumina Hiseq. Metagenome reads were quality trimmed and filtered using rqcfilter tool from bbtools package (http://jgi.doe.gov/data-and-tools/bbtools/), which performs primer and adapter removal, trims reads to the quality of 10, and removes PhiX and human sequences. The resulting reads were error-corrected using BFC tool (https://github.com/lh3/bfc.git) (Li, 2015) with kmer length of 25 and removing reads containing unique kmers. The resulting filtered and error-corrected reads were assembled for each sample separately using SPAdes v.3.9.0 without error correction with kmers 27, 47, 67, 87, 107 (Bankevich et al., 2012). Reads were mapped to the assemblies using Burrows-Wheel Aligner (BWA) v0.7.15 (Li and Durbin, 2010) and binned based on abundance patterns and kmer composition using Metabat v0.32.4 with minimum contig length of 3 kb and superspecific probability option (Kang et al., 2015). Differential coverage could not be utilized as there was little overlap between the 3 OV-2 samples (i.e. less than 10% of the reads from P1 and P2 could be mapped to P3, and vice versa). The bins corresponding to Solagigasbacteria were identified based on the presence of Solagigasbacteria 16S rRNA on contigs longer than 20 kb, as well as best BLAST hits to Solagigasbacteria SAG assemblies. Additional Solagigasbacteria metagenome bins were identified by BLASTing annotated gene regions of SAGs against metagenome assemblies, and bins with ≥ 200 hits with ≤ 1e-50 evalue score were further analyzed with checkM v1.0.6. Bins with excessive contamination were ignored. Metagenome assemblies are deposited in the Integrated Microbial Genomes database at the Joint Genome Institute (3300009626, 3300009691, 3300009444 and 3300003659).

Recruitment of metagenome reads to single-amplified genomes (SAGs) was determined using in-house software and Burrows-Wheel Aligner (BWA) v0.7.15 (Li and Durbin, 2010) to map sequence reads to Solagigasbacteria SAG contigs that met the criteria of 100 bps overlap at ≥90 % nucleotide identity. The relative abundance of SAG relatives was determined as the fraction of metagenome reads mapping per megabase of a reference genome.

## RESULTS AND DISCUSSION

### 16S rRNA gene phylogeny and biogeography

We used full-length 16S rRNA gene sequences of Solagigasbacteria SAGs as queries in BLASTn searches for related sequences in the NCBI nucleotide database that yielded 91 unique sequences ≥ 85% nucleotide identity over ≥ 600 bps. A phylogenetic analysis of these sequences suggested that Solagigasbacteria form a strongly bootstrap-supported, monophyletic lineage (Figure 1). Nitrospirae was the most closely related phylum, sharing 79 – 83% 16S rRNA gene sequence identity with Solagigasbacteria. Some Solagigasbacteria 16S rRNA gene sequences were misclassified as Nitrospirae in public databases (green arrows in Figure 1, also see Supplemental Figure 3). Given that the Solagigasbacteria are a bootstrap-supported, monophyletic clade, contain unifying genomic features (e.g. GC content), and fall below the median phylum-level 16S rRNA gene similarity threshold of 83.68% (range 81.6-85.93%) (Yarza et al., 2014), we propose that Solagigasbacteria is a unique phylum-level lineage. Phylogenies based on ribosomal preotein sequences (Hug et al., 2016a) (Anantharaman et al., 2016) agree with this phylogenetic placement and phylum demarcation. A superphylum may be formed by Solagigasbacteria, Aminicenantes, Acidobacteria, and the Candidate phylum NC10, but further phylogenomic analyses are needed to confirm this hypothesis.

**Figure 3.**
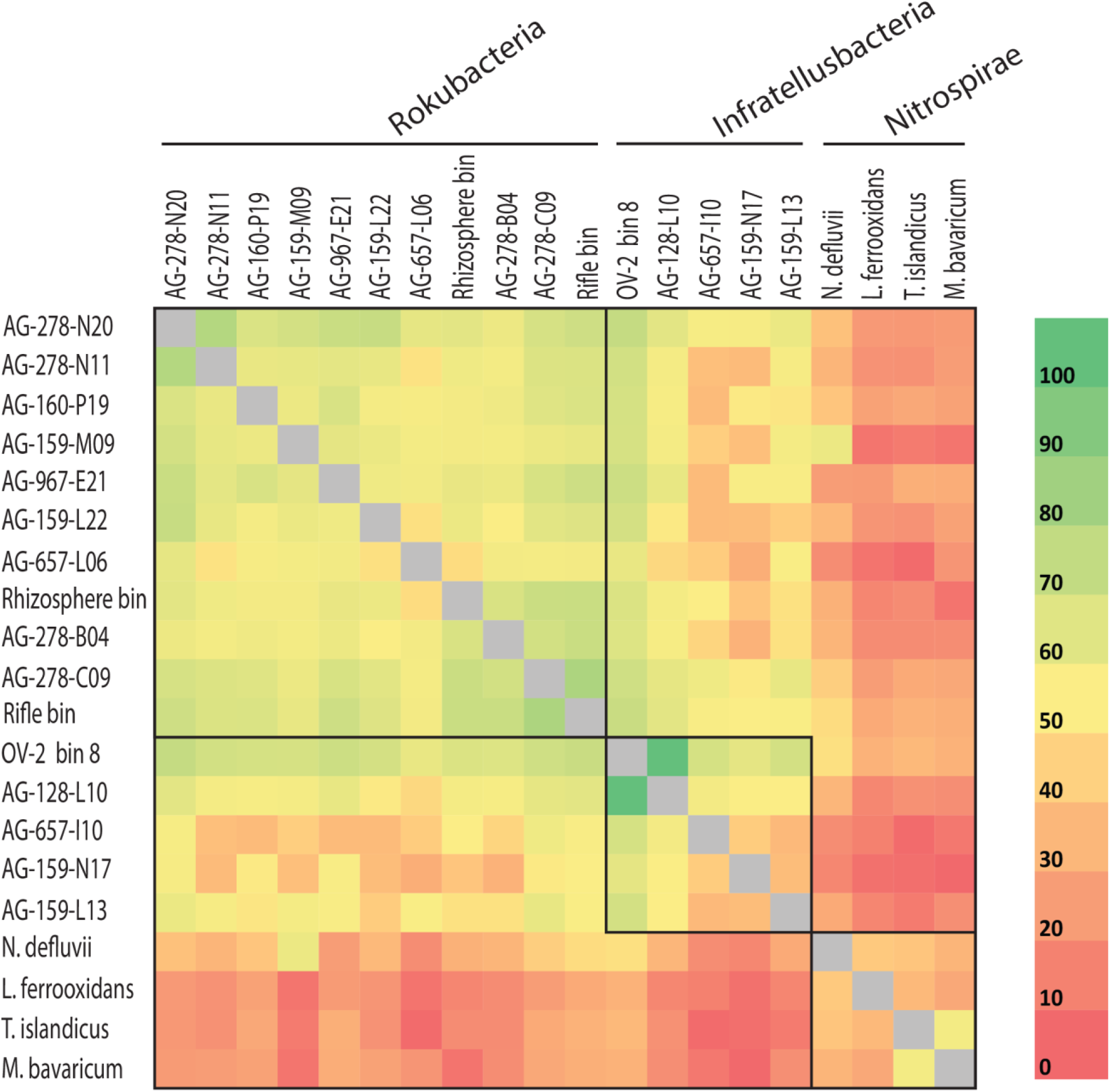
Average amino acid identity (AAI) for shared proteins among more complete Solagigasbacteria single amplified genomes (SAGs) (>0.5 Mbps) and Solagigasbacteria metagenome bin8 and rhizosphere bin (indicated by asterisks in Table 1), and select genomes from the Nitrospirae phylum. Boxes indicate phylogenetically defined class-level lineages or above (also see Figure 1 and Supplemental Table 2).

The Solagigasbacteria 16S rRNA gene sequences form two deeply branching sub-lineages that diverge from each other by ∼12 – 15%, i.e. at an operationally-defined class level (Figure 1; Supplemental Table 2) (Hugenholtz et al., 1998; Yarza et al., 2014). Apart from SAGs and PCR-derived sequences, one of the sub-lineages also included 16S rRNA genes from metagenome bins obtained from the Puerto Rican *Tabebuia* rhizosphere (light blue square in Figure 1) and from a previously published bin from the Rifle site, Colorado (orange square in Figure 1) (Anantharaman et al., 2016). The latter study named this lineage candidate phylum Rokubacteria, and we propose retaining this name for a class within Solagigasbacteria. Rokubacteria encompassed the majority of Solagigasbacteria sequences originating from both soils and shallow terrestrial subsurface environments. The majority of Rokubacteria SAGs from OV-2 fell into a subclade comprised exclusively of 16S rRNA gene sequences from subsurface sites. The second class-level lineage included a smaller set of sequences that originate exclusively from terrestrial subsurface sites. We propose naming this candidate class “Candidatus Infratellusbacteria” (hereby referred to as Infratellusbacteria), in order to reflect the predominant environment in which these microorganisms have been detected so far.

The sources of samples from which Solagigasbacteria 16S sequences were retrieved (25 in total; including 19 previously sampled sites (Figure 1), suggest a cosmopolitan distribution in soils and terrestrial subsurface, with no evidence so far for presence in marine environments. Interestingly, Solagigasbacteria were low in abundance at almost every site where they were identified in this and prior studies (Lin et al., 2012; Figuerola et al., 2015), and often were represented by a single 16S rRNA gene sequence. An alternative analysis of Solagigasbacteria abundance in our study sites, by performing metagenome fragment recruitment on SAGs as references, provided further evidence that Solagigasbacteria comprised ∼1 % of the microbial community in OV-2 (Supplemental Figure 4), similar to other samples (Lipson, 2004; Hansel et al., 2008; Chen et al., 2012; Lin et al., 2012; Rodrigues et al., 2014; Figuerola et al., 2015). A recent study identified Rokubacteria to constitute ∼10 % of the microbial community in a grass root zone in the Angelo Coast Range Reserve, California, making it the most Solagigasbacteria-rich environment to date (Butterfield et al., 2016), though no 16S rRNA sequences were identified in the metagenome bins.

**Figure 4.**
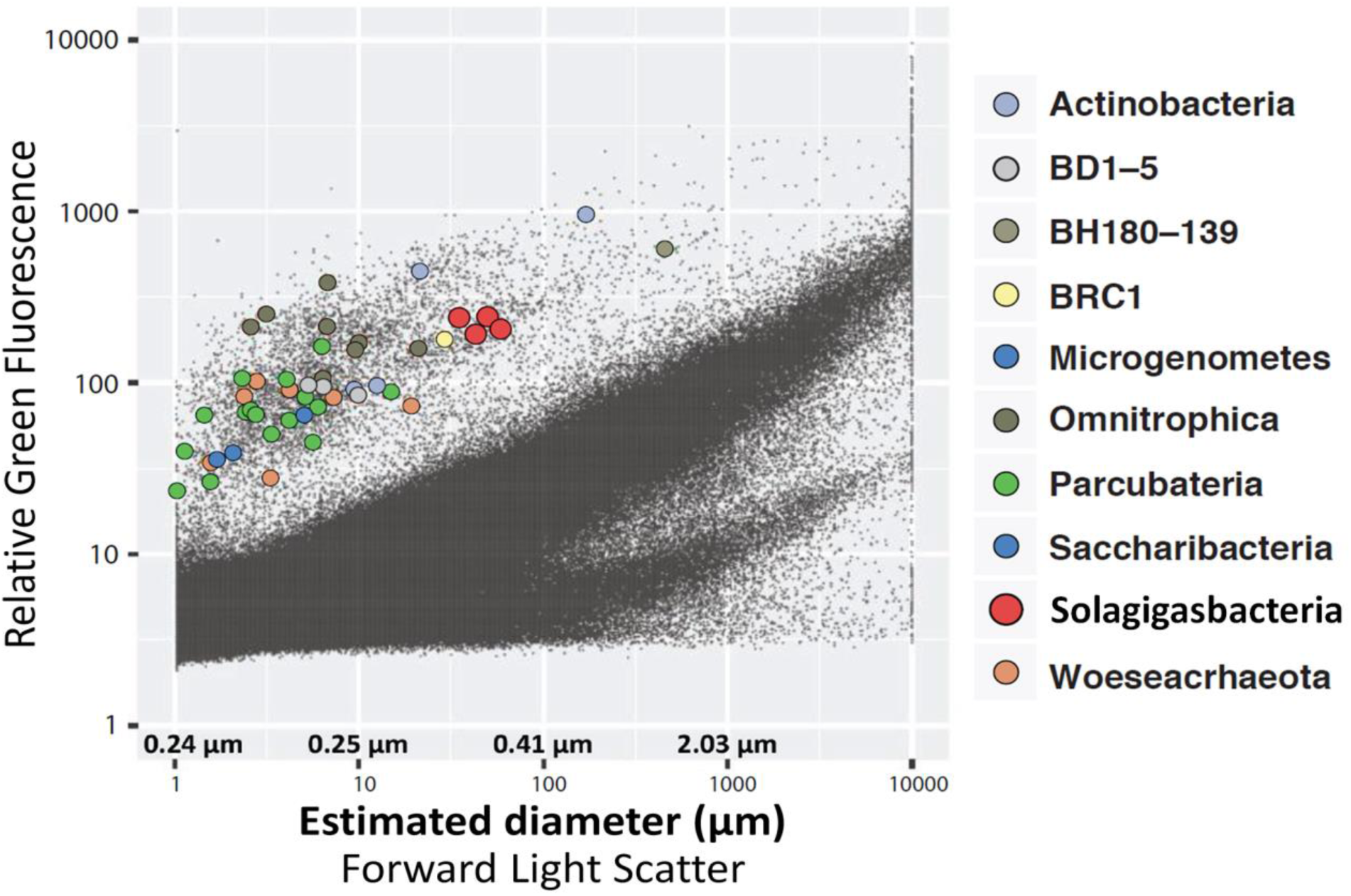
Optical properties and estimated diameters of cells sorted from the OV-2 sample that contained the largest number of cells identified as Solagigasbacteria. Colored dots indicate cells that were successfully identified by their 16S rRNA gene. Black dots indicate all particles detected by the fluorescence-activated cell sorter.

### General genome features

The SAGs obtained from SURF, Finsch, OV-2 and Crystal Spring sites contained phylogenetically diverse representatives of both Solagigasbacteria classes, enabling us to explore their genomic content, metabolic potential and evolutionary histories. *De novo* genome assemblies of the 19 SAGs ranged from 0.05 to 2.86 Mbps (Table 1). The estimated Solagigasbacteria genome completeness ranged between 1 – 40 % (average of 18.2 %). This is significantly lower than the genome recovery from other SAGs using the same techniques in earlier studies, which averaged at around 50 % (Rinke et al., 2013; Swan et al., 2013; Kashtan et al., 2014). Based on the presence of conserved single copy genes in the most complete SAG assemblies, we estimate that Solagigasbacteria complete genomes are 6 – 8 Mbps in length (average 6.8 Mbps; Table 1 and Supplemental Figure 5), which is slightly larger than estimates obtained from metagenome bins at the Rifle site (4 – 6 Mbps; Supplemental Table 3) (Anantharaman et al., 2016) and the Puerto Rican soil (Table 1). Genome size predictions for smaller SAGs and contaminated metagenome bins are variable, though the more complete SAGs and metagenome bins converge on the average genome size reported above (Supplemental Figure 5). A relatively large fraction, between 8 – 17 % of the Solagigasbacteria genomes, consists of nucleotides predicted to be non-coding. With a few intriguing exceptions (Sekiguchi et al., 2015), these features present a stark contrast to the predominantly small and streamlined genomes of most recently described bacterial and archaeal candidate phyla from diverse surface and subsurface environments, including the abundant and diverse candidate superphylum Patescibacteria (Rinke et al., 2013), which was later proposed to constitute an even larger evolutionary unit, the Candidate Phyla Radiation (CPR) (Rinke et al., 2013; Brown et al., 2015; Castelle et al., 2015; Anantharaman et al., 2016).

**Figure 5.**
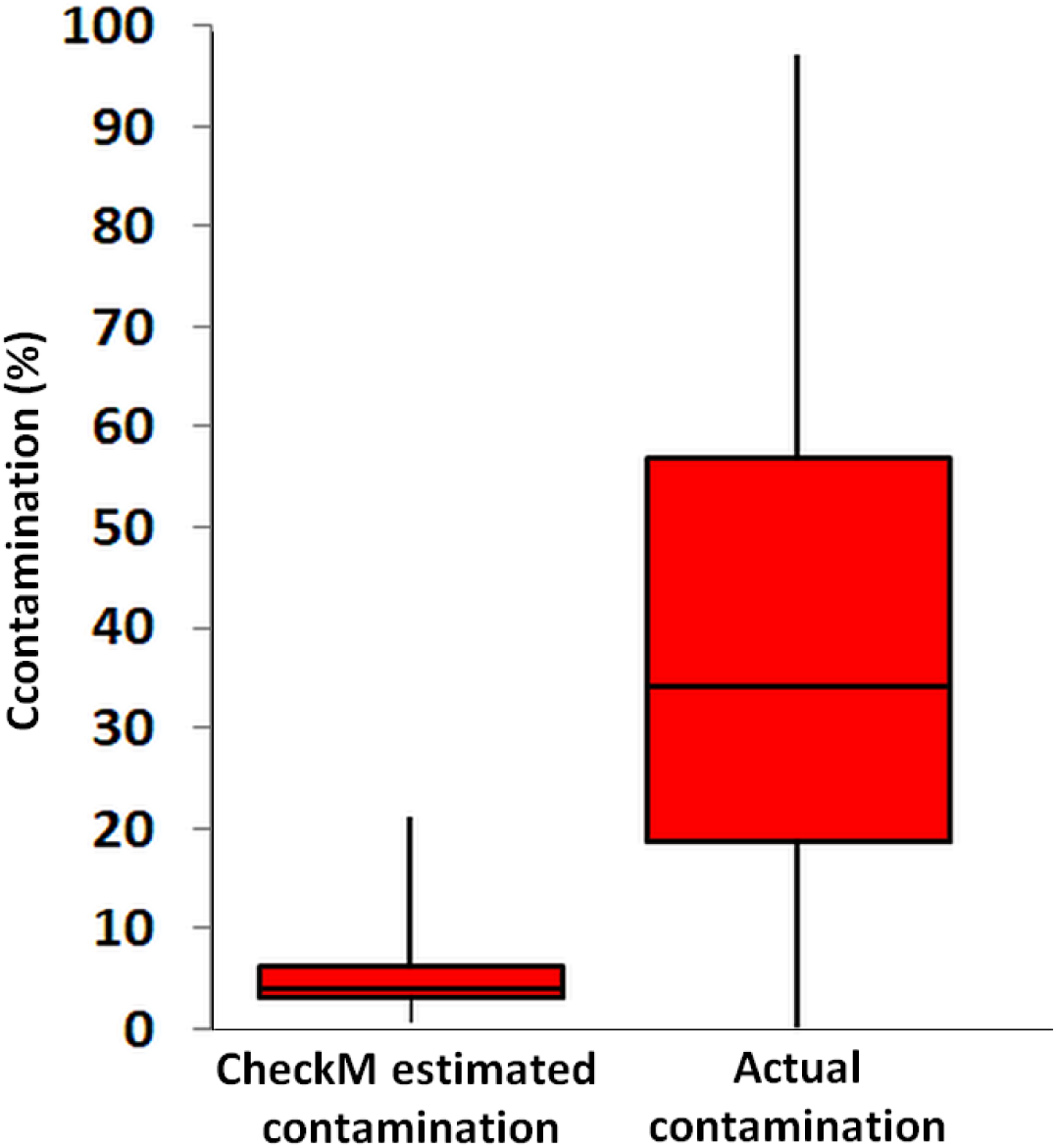
Contamination predicted by checkM software from all pairwise SAG combinations across the class-level lineages Rokubacteria and Infratellusbacteria (15 Rokubacteria and 4 Infratellusbacteria SAGs; 60 combinations total), compared to actual contamination calculated from all artificially combined SAGs. Two-sample t-test assuming equal variances was significant (p=<000.1).

**Table 1.** Solagigasbacteria assembly statistics and predicted genome completeness. Asterisks indicate more complete genome assemblies that were used in Figure 3.

| Solagigasbacteria<br>SAGs | Class | Site | Raw PE<br>reads<br>(millions) | Assembly<br>(Mbps) | GC<br>% | 16S<br>rRNA+ | Estimated<br>Genome<br>Completeness,<br>% | Predicted<br>genome<br>size-<br>Mbps | %<br>Contami-<br>nation<sup>4</sup> | Genome<br>quality<sup>5</sup> |
| --- | --- | --- | --- | --- | --- | --- | --- | --- | --- | --- |
| **\*AD-967-E21** | Rokubacteria | Finsch | 13.7 | 1.52 | 66 | Yes | 30 | 5.1 | <1 | low |
| **\*AG-128-L10** | Infratellusbacteria | CS<sup>1</sup> | 13.2 | 1.89 | 70 | Yes | 24 | 7.9 | <1 | low |
| AG-138-D06 | Rokubacteria | SURF<sup>2</sup> | 12.0 | 0.29 | 67 | Yes | 6 | 4.9 | 0 | low |
| AG-657-A01 | Rokubacteria | OV-2 | 12.2 | 0.40 | 68 | Yes | <1 | NA | 0 | low |
| **\*AG-657-I10** | Infratellusbacteria | OV-2 | 10.8 | 0.93 | 71 | Yes | <1 | NA | 0 | low |
| **\*AG-657-L06** | Rokubacteria | OV-2 | 7.7 | 1.03 | 65 | Yes | <1 | NA | 0 | low |
| AG-159-B15 | Rokubacteria | OV-2 | 8.8 | 0.05 | 64 | Yes | <1 | NA | 0 | low |
| AG-159-G23 | Rokubacteria | OV-2 | 7.6 | 0.40 | 64 | No | 5 | 8.1 | 0 | low |
| **\*AG-159-L22** | Rokubacteria | OV-2 | 8.4 | 0.69 | 65 | Yes | 16 | 4.3 | 0 | low |
| **\*AG-159-M09** | Rokubacteria | OV-2 | 7.5 | 0.93 | 65 | Yes | 4 | 22.2 | 0 | low |
| AG-159-N17 | Infratellusbacteria | OV-2 | 9.8 | 0.41 | 67 | Yes | <1 | NA | 0 | low |
| AG-159-P01 | Rokubacteria | OV-2 | 5.1 | 0.07 | 65 | No | <1 | NA | 0 | low |
| AG-160-L13 | Infratellusbacteria | OV-2 | 7.0 | 0.41 | 64 | Yes | 3 | 13.7 | 0 | low |
| **\*AG-160-P19** | Rokubacteria | OV-2 | 8.2 | 1.38 | 64 | Yes | 27 | 5.1 | <1 | low |
| AG-278-A09 | Rokubacteria | OV-2 | 0.08 | 0.24 | 65 | Yes | 4 | 6.1 | 0 | low |
| **\*AG-278-B04** | Rokubacteria | OV-2 | 8.4 | 2.61 | 69 | Yes | 32 | 8.2 | <1 | low |
| **\*AG-278-C09** | Rokubacteria | OV-2 | 8.5 | 1.68 | 68 | Yes | 18 | 9.21 | 0 | low |
| **\*AG-278-N11** | Rokubacteria | OV-2 | 7.4 | 1.15 | 67 | Yes | 15 | 7.8 | 0 | low |
| **\*AG-278-N20** | Rokubacteria | OV-2 | 0.16 | 2.86 | 67 | Yes | 40 | 7.2 | 0 | low |
| **Solagigasbacteria**<br>**metagenome bins** |  |  |  |  |  |  |  |  |  |  |
| **\*OV-2 bin8<sup>3</sup>** | Infratellusbacteria | OV-2 | - | 5.69 | 72 | yes | 89 | 6.4 | 2 | Medium |
| **\*Rhizosphere bin<sup>3</sup>** | Rokubacteria | PR | - | 4.01 | 69 | yes | 63 | 6.4 | 1 | Low |
| OV-2 bin1 | unknown | OV-2 | - | 11.84 | 62 | no | 100 | 11.8 | 332 | NA |
| OV-2 bin2 | unknown | OV-2 | - | 11.68 | 69 | no | 100 | 11.6 | 175 | NA |
| OV-2 bin6 | unknown | OV-2 | - | 6.06 | 69 | no | 73 | 8.3 | 59 | NA |
| OV-2 bin9 | unknown | OV-2 | - | 5.05 | 71 | no | 78 | 6.5 | 34 | NA |
| OV-2 bin11 | unknown | OV-2 | - | 4.68 | 68 | no | 98 | 4.8 | 153 | NA |
| OV-2 bin43 | unknown | OV-2 | - | 1.32 | 72 | no | 14 | 9.5 | 1 | Low |

<sup>1,2</sup> Crystal Spring, Nevada and Sanford Underground Research Facility (300 m).
<sup>3</sup> Metagenome bins from OV-2 and *Tabebuia* rhizosphere sample taken in Puerto Rico.
<sup>4</sup> Estimated with checkM
<sup>5</sup> Genome quality reported according to Bowers et al., 2017.

The GC content of Solagigasbacteria SAG assemblies was at the high end of the reported spectrum for known organisms, ranging between 64 – 71 %, with an average of 68% (Table 1; Figure 2). This is in agreement with the high % GC content of the Rokubacteria metagenome bins reported by Anantharaman et al. (Anantharaman et al., 2016). The most closely related phylum to Solagigasbacteria, Nitrospirae, has a more variable GC content, ranging from 34 % (*Thermodesulfovibrio islandicus*) to 62% (*Nitrospira moscoviensis*). The factors determining GC content remain unclear. The spontaneous mutations may favor nucleotide shifts to A and T (Hershberg and Petrov, 2010; Hildebrand et al., 2010), and the lower nitrogen content of AT may provide a selective advantage to low GC organisms in N-limited environments (Giovannoni et al., 2014). Yet, high %GC is present in a wide range of lineages and habitats (Hershberg and Petrov, 2010). Factors determining high %GC remain controversial, with some studies suggesting the importance of temperature and solar radiation as selective variables (Foerstner et al., 2005; Hildebrand et al., 2010), while other reports refute these findings (Lassalle et al., 2015; Li et al., 2015). Furthermore, while some studies suggest GC content is evolutionarily conserved within lineages (Lassalle et al., 2015; Reichenberger et al., 2015), other studies show large GC variation among lineages that were thought to be exclusively high in GC, such as the Actinobacteria phylum (Ghai et al., 2012; Swan et al., 2013) and the Roseobacter clade of Alphaproteobacteria (Swan et al., 2013; Zhang et al., 2016). The high %GC of Solagigasbacteria contrasts low %GC in most of the major, uncultured branches of Bacteria and Archaea explored with single-cell genomics (Rinke et al., 2013) and metagenome binning (Anantharaman et al., 2016; Hug et al., 2016b) to date (Figure 2). It remains to be understood what evolutionary processes are involved in the emergence and maintenance of high %GC, and to what extent has the discovery of novel microbial lineages with high %GC been hampered by biases in DNA amplification (Stepanauskas et al., 2017) and sequencing techniques (Chen et al., 2013).

Solagigasbacteria SAG assemblies shared between 36 and 922 orthologous protein-encoding genes (average of 308 reciprocal orthologous protein hits). The average amino acid identity (AAI) was 46.2% (range from 34.6 – 64.2%; Figure 3), demonstrating high cell-to-cell genome divergence. Interestingly, SAGs originating from the OV-2 sample shared roughly the same proportion of protein-coding genes as SAGs from geographically distant sites. The most divergent Solagigasbacteria SAGs were obtained from the same OV-2 site (Figure 1), both within and between class-level lineages. Genomes were mostly non-syntenic on larger scales. However, many shared proteins of related function were located in small islands of synteny in the six least fragmented SAG assemblies (Supplemental Figure 6). Causes for the unusually variable genome content among cells in each study site remain unclear. Dispersal of dormant cells to the sampling sites from a multitude of evolutionarily distant populations is one plausible explanation. An alternative explanation may be the accumulation of point mutations, gene acquisitions, gene loss and genome rearrangements at a rate that outpaces cell division. The latter possibility is highly speculative and contradicts conventional models of microbial evolution, but should be viewed in the context of bacterial generation times potentially ranging in hundreds and even thousands of years in some low-energy, subsurface environments (Labonte et al., 2015a).

Solagigasbacteria genomes contain numerous transposases and integrases (4 – 60 per SAG assembly; Supplemental Table 4). Genes of potential viral origin and CRISPR regions were also identified in most Solagigasbacteria SAGs (Supplemental Table 4). The contigs that contained phage-like genes were never predicted to be entirely viral, indicating integration into host chromosomes. These observations are similar to the recent finding of abundant transposable prophages in Firmicutes in the deep subsurface of the Witwatersrand Basin (Labonte et al., 2015a) and indicate a potentially important role of viruses as vectors of horizontal gene transfer in low-energy, subsurface environments.

### Predicted phenotype and energy production

We employed forward light scatter (FSC) signals from FACS, which where calibrated against a series of benchmark cultures, to estimate approximate diameters of the cells from which SAGs were generated (Stepanauskas et al., 2017). This indicated that Solagigasbacteria cell diameters ranged between 0.3 – 0.4 μm (Figure 4). While this estimate is greater than the 0.15 – 0.20 μm diameter reported for some of the CPR cells (Luef et al., 2015), and the ∼0.2 –0.3 μm average diameter of the most abundant marine bacterioplankton lineage *Pelagibacter* (Giovannoni et al., 2005; Giovannoni et al., 2014), it is approaching the theoretical lower limit for cell sizes (NRC., 1999). Such small cells, including the CPR, *Pelagibacter, Mycoplasma*, ultrasmall Actinomycetes, and *Prochlorococcus*, have extremely small, streamlined genomes that range between 0.8 – 2.5 Mbps (Biller et al., 2014; Nakai et al., 2016; Parrott et al., 2016). In the case of Solagigasbacteria, the presence of large genomes in small cells may imply extensive DNA packaging or dormancy. In partial support of this hypothesis, a variety of DNA packaging and super-coiling proteins where annotated in the Solagigasbacteria SAGs and metagenome bins (Supplementary Table 5). Further experimental work is required to confirm these predictions.

Solagigasbacteria contain numerous genes that are typical of gram-negative (diderm) organisms, including the majority of genes involved in the production and transport of lipids across the cytoplasmic membrane for outer membrane and LPS assembly (Sutcliffe, 2010) (Supplemental Figure 1 and Supplemental Table 5), which is consistent with their phylogenetic affiliation with the Gram-negative Nitrospirae. We identified multiple genes involved in twitching motility in 11 Rokubacteria SAGs, 4 Infratellusbacteria SAGs, and both metagenome bins, possibly indicating a conserved mechanism of pili motility in the Solagigasbacteria (Supplemental Table 5). We also identified genes in 3 Rokubacteria SAGs and Infratellusbacteria OV-2 bin 8 that are predicted to encode flagella structural proteins, while propeller filament genes were absent in all SAG assemblies and metagenome bins (Supplemental Figure 1). While OV-2 bin 8 contained genes involved in flagella assembly, Infratellusbacteria SAGs lacked genes required for the assembly of flagella, though gene absence could be due to fewer and less complete SAG assemblies. Furthermore, putative genes were identified in the majority of assemblies for methyl-accepting chemotaxis proteins, two-component sensor kinases, ATP motor proteins, and sensor proteins for nitrogen, oxygen, zinc/lead, and acetoacetate, indicating that Solagigasbacteria can respond to a broad range of chemical stimuli. Solagigasbacteria contain multiple carbon transport proteins, including those specializing in lipids, peptides and sugars, and carbon degradation and election transport pathways for aerobic respiration (Supplemental Figure 1). Rokubacteria SAGs also contain genes involved in nitrogen respiration, including nitrite oxidoreductases, which are universally conserved in the Nitrospirae lineage (Supplemental Figure 7). See Supplemental Section I for detailed metabolic predictions.

### Comparison of Solagigasbacteria assemblies from single cells and metagenomes

The availability of several partial genomes of Solagigasbacteria from single cells and metagenomes from this and prior studies (Table 1 and Supplemental Table 3) offered an opportunity to compare the type and quality of information that can be extracted using these two approaches. Genome completeness is one important quality metric of *de novo* assemblies. The ratio of the number of single copy marker genes that are found versus expected in an assembly is the most commonly use proxy for genome completeness and is implemented in popular computational tools, such as checkM (Parks et al., 2015). In our study, checkM-based estimates of assembly completeness of Solagigasbacteria SAGs and metagenome bins ranged between 1-40% and 14-100%, respectively, suggesting that individual SAG assemblies tended to be less complete than metagenome bins (Table 1). However, the higher average completeness of metagenome bins came with important caveats: high estimated contamination in five out of eight bins (Table 1), absence of rRNA genes in six out of eight bins, and the lack of knowledge of the number and genetic diversity of cells that contributed genomic sequences to each bin. These caveats may limit the interpretability of metagenome bins in the context of microbial ecology and evolution and outweigh the benefits of their higher estimated completeness.

While the checkM-based estimates of SAG contamination were always below 1%, they ranged between 1-332% (average 95%) in our OV2 for metagenome bins (Table 1) and between 2-14% in bins of an earlier study (Anantharaman et al., 2016) (Supplemental Table 3), suggesting quality limitations of most bins (Table 1). These observations are in general agreement with the recent benchmarking effort employing > 1,000 previously sequenced strains of microorganisms and mobile genetic elements, which found that the performance of metagenome assembly and binning is impaired by the presence of related strains in a sample (Sczyrba, 2017).

The checkM estimates of contamination are based on the phylogenetic placement of the assembly’s single copy marker genes against a built-in database (Parks et al., 2015), which lacks many uncultured lineages, including Solagigasbacteria. To the best of our knowledge, the ability of checkM to detect contamination that originates from lineages that are absent from its database has never been evaluated. To address this question, we created pairwise combinations of assemblies of each Infratellusbacteria SAG with each Rokubacteria SAG. The checkM-estimated contamination in these combined assemblies was significantly smaller than the real, cross-class contamination (Figure 5), suggesting that checkM may fail detecting contamination from lineages not represented in the checkM database. Strikingly, the majority of our artificial combinations of SAGs from different phylogenetic classes would be considered ‘high quality’ genomes according to recently proposed genome standards for SAGs and metagenome bins (Bowers, 2017). In order to assess whether similar, cross-class contamination may be affecting our metagenome bins, we analyzed AAI among Solagigasbacteria SAGs and the only OV2 metagenome bin that contained a rRNA gene (bin8). While the rRNA gene placed this bin firmly among the Infratellusbacteria (Figure 1), its AAI suggested affiliation with Rokubacteria (Figure 3). Furthermore, the best BLAST hits to bin8 genes consisted of SAGs from both class-level lineages, including multiple near full-length alignments at > 95% nucleotide identity with Rokubacteria SAGs (Supplemental Table 6). This indicates that the checkM-based estimate of 2% contamination for this bin may be a major underestimate. These findings imply that improvements are urgently needed in the quality control of genome assemblies originating from uncultured microbial groups and in the validation of the performance of QC software.

The comparison of SAGs and metagenome bins demonstrates that the two approaches provide two fundamentally different types of data and should be interpreted accordingly. While SAG assemblies represent fragments of discrete genomes from individual cells, the metagenome bins are fragments of consensus sequences derived from a multitude of genetically non-identical organisms. The consistency of certain general features between Solagigasbacteria SAGs and bins (e.g. high %GC, large estimated genome size, and many shared metabolic pathways) suggests that metagenome bins provide useful consensus information about this candidate phylum (Figure 2 and Supplemental Figure 1). However, consensus sequences appear to mask extensive genetic diversity among Solagigasbacteria cells in the studied environments. On a more fundamental level, metagenome assembly and binning relies on the assumption that microbial communities are composed of near-clonal populations. An increasing body of evidence shows that this assumption is not valid in many microorganismal lineages and environments, with genomic rearrangements and horizontal gene transfer being more prevalent than previously thought (Ochman et al., 2000; Feldgarden et al., 2003; Shapiro, 2010; Kashtan et al., 2014; Labonte et al., 2015a). By recovering data from the most fundamental units of biological organization, single-cell genomics does not rely on the assumption of clonality, offers an opportunity to improve our understanding of microbial microevolutionary processes (Garrity and Lyons, 2003; Engel et al., 2014; Kashtan et al., 2014), and helps calibrating the performance and interpretation of metagenomics tools when working with complex, natural microbial assemblages (Becraft et al., 2015).

## CONCLUDING REMARKS

Recent discoveries of many novel phyla and superphyla of microorganisms are revolutionizing our understanding of the genealogy and current diversity of life. Here, a focused analysis of the single cell genomic and metagenome sequences of Solagigasbacteria (formerly known as SPAM) suggests that they constitute a monophyletic, phylum-level lineage that is most closely related to Nitrospirae among the currently described phyla. Large genomes, high %GC, and a global presence at low abundance in soils and terrestrial subsurface environments appear to be general features of this candidate phylum. Solagigasbacteria genomes predict didermy, mixotrophy, motility, and versatile DNA packaging mechanisms. It is plausible that the latter feature interferes with gDNA amplification, in part explaining the difficulty of recovering high quality genomes from Solagigasbacteria single cells. Furthermore, large cell-to-cell genomic heterogenetity and low relative abundance in most environments studied to date may be among the factors contributing to their limited recovery in metagenome bins. Our analysis also demonstrates major differences in the quality of genomic data obtained from SAGs and metagenome bins: While assemblies with greatest estimated genome recovery where obtained by metagenome binning, SAGs delivered contamination-free data from discrete biological units, making them easier to interpret and revealing significant genomic diversity within this candidate phylum, including a split into two class-level lineages.

## ACKNOWLEDGEMENTS

We thank the staff of the Bigelow Laboratory Single Cell Genomics Center and the Joint Genome Institute for the generation of single cell and metagenomic data. We are grateful to Olukayode Kuloyo, Borja Linage, Sarah Hendrickson, Cara Magnabosco, Melody Lindsay, Maggie Lau, Petra Diamonds and the management and staff for Finsch diamond mine, South Africa for their help obtaining the samples. We are also grateful to Laura Vann for their help obtaining Nevada field samples. This work was supported by the U.S. National Science Foundation grants DEB-1441717 and OCE-1335810. The work conducted by the U.S. Department of Energy Joint Genome Institute is supported by the Office of Science of the U.S. Department of Energy under Contract No. DE-AC02-05CH11231. **The authors declare that there are no conflicts of interest.**

